# Male mouse strain variation reveals divergent phenotypes for extrinsic and intrinsic reward motivation

**DOI:** 10.64898/2026.08.11.743966

**Authors:** Eleanor W Grayson, Emma S J Robinson, Megan G Jackson

**Affiliations:** School of Physiology, Pharmacology and Neuroscience, Biomedical Sciences Building, University of Bristol, Bristol, England

## Abstract

Motivational deficit is a prevalent symptom across a wide range of neurodegenerative and neuropsychiatric disorders. Despite its clinical importance, first-line treatments for these disorders fail to effectively treat this symptom domain. In animal models, motivation is typically assessed in the context of extrinsic reward, where reward is delivered for completing an effortful action. However, many motivated behaviours occur in the absence of a tangible reward and are instead driven by intrinsic motivation. Previous work has shown that an extrinsic motivation task (effort for reward (EfR)) and an intrinsic motivation task (effort based forage (EBF) task) show opposing responses to a range of pharmacological manipulations. However, it is not clear whether intrinsic and extrinsic motivation dissociate in the context of endogenous behavioural variation. We therefore investigated whether these tasks were sensitive to behavioural variation across three different strains of mice (C57Bl/6JJRi, 129S2/SvPasOrlRj and BALB/cJRi) and whether strain profiles diverged across tasks. Here, we found that BALB/c mice showed the lowest levels of foraging in the EBF task, indicative of a low intrinsic motivational state but showed the highest levels of high effort responding in the EfR task, indicative of a high extrinsic motivational state. These differences were not driven by an anxiety-related phenotype and were therefore indicative of a motivation phenotype divergence across tasks. This work highlights the importance of moving away from considering motivation on a single axis, as findings can diverge depending on the nature of the motivational process. This has important implications for both phenotypic interpretation and the development of treatments targeting motivational dysfunction.

---

Loss of motivation is a transdiagnostic symptom domain with high prevalence across multiple psychiatric and neurodegenerative disorders (Epstein and Silbersweig, 2016). It has a significant impact on patient quality of life and is associated with greater functional impairment and cognitive decline (Fervaha et al., 2016, Forstmeier and Maercker, 2015). First-line treatments for these disorders including widely used antidepressants, antipsychotics and cognitive enhancers fail to effectively treat motivational deficit and, in some cases, can even worsen symptoms (Sansone and Sansone, 2010, Fervaha et al., 2015, Rea et al., 2014). Therefore, motivational deficits represent a clinically important yet insufficiently addressed treatment target.

In animal models, motivation is typically assessed in the context of extrinsic reinforcement. In operant conditioning paradigms, specific instrumental behaviours such as nose pokes or lever presses are shaped over time by the delivery of a palatable food reward. The amount or the vigour with which an animal completes the conditioned action provides a readout of motivational state. The Effort for Reward (EfR) task is based on these principles with the addition of a decision-making component in which the rodent must choose between making an effortful number of nose pokes to receive palatable reward or consume freely available standard laboratory chow. The allocation of behaviour to each option provides an operationalised measure of motivated behaviour (Salamone et al., 1991, Marangoni et al., 2023). Tasks such as these have been vital in revealing a role for the dopaminergic mesolimbic system in motivated behaviour (Salamone et al., 2007, Cousins et al., 1996) thereby positioning it as a target for therapeutics and a potential pathological mechanism in motivation disruption (Le Heron et al., 2018a, Muhammed et al., 2016).

However, it is important to recognise that many clinically relevant experiences of motivational impairment, such as reduced engagement in hobbies, exploration, or self-initiated activities termed ‘activities of daily life’ (ADL), often occur in the absence of explicit external reinforcement and are instead driven by intrinsically motivated processes. In this domain of motivation, the behavioural action itself holds reward value and is not contingent on a clear external reward. Consequently, tasks relying exclusively on extrinsic reward may capture only part of the motivational dysfunction observed in psychiatric disorders (Morris et al., 2022).

The Effort Based Forage task provides a unique opportunity to tap into intrinsically motivated processes. In this task, the mouse is provided with food and water and can freely choose to leave a safe and enclosed environment, and forage for nesting material in a more open environment, which requires a degree of effort to obtain (Grayson et al., 2025). The mouse shows similar rates of foraging when nesting material is freely available and when the environment is warmed, suggesting the act of foraging itself is in part intrinsically rewarding (Xeni et al., 2024).

While both are considered at their broadest level, measures of motivation, the Effort for Reward task and Effort Based Forage task show opposing responses to a range of serotonergic antidepressant compounds but remain broadly similar in the context of dopaminergic manipulation (Xeni et al., 2024, Xeni et al., 2025). Differences in behavioural response to class-specific compounds suggests that the two domains of motivation may be driven by partially distinct neural circuits. However, while acute pharmacological manipulation provides insight into mechanism via transient modulation of neurochemical systems, it remains unclear whether intrinsic and extrinsic motivation dissociate in the context of endogenous behavioural variation. Dissociation at this level would provide further evidence that these tasks capture distinct motivational phenotypes rather than task-specific responses to pharmacological manipulation.

Behavioural variation is particularly evident between strains of mice, which show differing profiles across a wide range of behavioural domains relevant to psychiatric disorders such as exploratory activity, sociability, anxiety-like behaviours and cognitive processing. For example, five commonly used strains of mice including C57Bl/6JJ, BALB/c, CBA, 129SvEv and CD1 showed distinct behavioural phenotypes when assessed using a battery of tests (Sultana et al., 2019). As such, strain variation provides an opportunity to determine whether intrinsic and extrinsic motivation show dissociable patterns of behavioural variation in the absence of overt pathology or transient pharmacological manipulation. In addition, motivation-related phenotypic differences between strains remain poorly characterised but have the potential to provide insight into neurobiological and treatment sensitivity differences underpinning distinct motivational phenotypes.

Therefore, in this study we identified three mouse strains commonly utilised in neuroscience research (C57BL/6JJRi, 129S2/SvPasOrlRj and BALB/cJRi) to investigate whether strain influences motivational phenotype in the EfR and EBF tasks. We hypothesised strain variation would differ across EfR and EBF tasks, consistent with these paradigms capturing partially distinct motivational domains.

## Methods

### Subjects

Two cohorts of 18 male mice comprising three different strains were used across all experiments (n=12 C57Bl/6JJ, n=12 129sv and n=12 BALB/c, Janvier, France, see **S1** for cohort information). Mice were aged 6 weeks on arrival and were singly housed in open-top Techniplast 1284 cages to mitigate aggressive behaviour (Davies et al., 2025) (see **S2** for further husbandry information). Mice were kept in a 12:12h reverse light cycle with lights off at 8:15am and on at 8:15pm. All experiments were conducted during the active phase with lights off, except anxiety-like behavioural assays which were conducted in white light. Holding rooms were temperature and humidity-controlled to ∼21°C, and mice were given standard laboratory chow (Purina, UK) and water *ad libitum* throughout testing except for the food motivated assays described below. Mice were habituated to cup handling before experiment onset in accordance with the 3Hs Initiative (3Hs, 2024) which began one week after arrival to allow for acclimatisation. All experiments were performed in accordance with the Animals Scientific Procedures Act (ASPA, UK) and approved by University of Bristol Animal Welfare and Ethical Review Body (AWERB). Behavioural tests were ordered (**S3**) to mitigate the impact of lasting behavioural effects on subsequent assays. For instance, operant training and EfR testing were performed last so that any long-lasting effects of food restriction or conditioned behaviour would not impact any following experiments. Mice were monitored for stereotypic behaviour throughout the study. Mice that showed harmful stereotypic behaviour were removed from the study. N = 5 mice were removed from the study. Exact details of which studies this affected are outlined below.

### Effort-based Forage task arena

The arena consists of an enclosed home area, an open forage area, a tube connecting these two areas, and a nesting material box secured in the forage area. The nesting material box is filled with white Sizzlenest (Datesand, UK) which mice can pull through apertures in the faceplate of the box (**fig. 1.AsB**). Dimensions of the EBF arena are provided in Supplementary **S4 s 5**. Alterations to the task, shown in **fig. 1.CsD**, can be applied to modulate effort needed to forage (aperture size of the faceplate) and produce an affective response to a more aversive area (size of the forage area). 3D renders of the task design and 3D print files for the nesting material box can be found here https://github.com/meganjackson13/Bedding-box-3D-files.

**Fig. 1.**
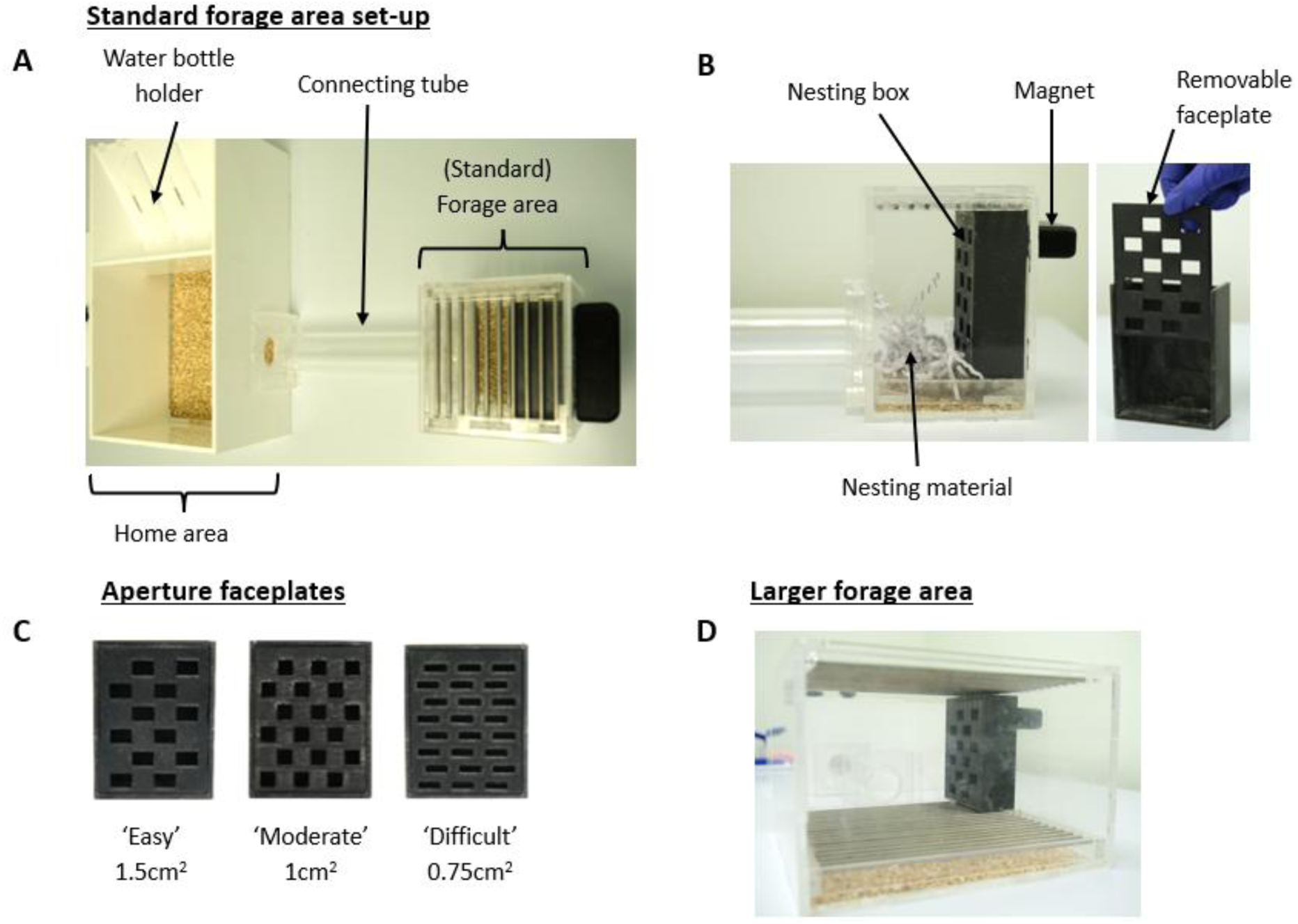
Effort Based Forage task arena. **A)** The home area is an enclosed area containing freely available standard lab chow (Purina, UK) and water and is connected via a tube to the open and therefore more aversive forage area containing the nesting box. Mice travel from the home box, through the connecting tube, to the forage area, ‘forage’ (pull out) nesting material from the nesting box with their mouth/paws, and ‘shuttle’ (carry back) the material to the home area. **B)** The nesting box is secured in the forage area by a magnet and filled with 18g of nesting material which is slightly pulled through the apertures of a faceplate so mice can reach it. The faceplate of the nesting box can be removed and changed. **C)** Three different sized aperture faceplates are used to alter effort required to forage during the effort curve paradigm, with a smaller aperture size requiring more effort needed to forage material. Habituation to the EBF arena uses the ‘Easy (1.5 cm^2^)’ aperture faceplate. **D)** The larger forage area is used to induce an affective response and observe how foraging behaviour changes with stress induced by the more aversive environment (affective reactivity test). The nesting box is secured immediately to the left of the tube (as the mouse enters the area).

### EBF task habituation

Mice underwent three initial sessions in the EBF arena without the nesting box present in the forage area. Mice were placed into the home area and left to explore for 10 minutes on the first day, then 5 minutes on the second and third days. To record the number of times the mouse entered the forage area as a measure of initial exploration, the experimenter remained in the room and tallied the number of times each mouse entered the forage area (with all four limbs crossed from the tube into the area). This was termed a ‘bout’ and was normalised to time (bouts/minute).

Next, the mice were left in the arena for a 4-hour session with the nesting box filled with 18g of nesting material, the ‘easy’ 1.5cm^2^ aperture faceplate in place and secured in the forage area. The mice had free access to standard lab chow and their water bottles in the home area for the entire session, and all sessions were run in darkness. Subsequently, three 2-hour baseline sessions were performed using the ‘moderate’ 1cm^2^ aperture faceplate. After each session the weight of the nesting box was measured as well as weight of nesting material left on the barred floor of the forage area (**Fig.2B**).

During these sessions, the mouse could freely choose to traverse through the arena and forage nesting material. Across all studies the main output measures were total material foraged (weight of nesting box before session – weight of nesting box after session) which reflects the amount of nesting material pulled from the box, total material shuttled (total foraged – weight left on forage area floor) which represents the amount of nesting material pulled from the nesting box and taken through the tube to the home area and percentage nesting material shuttled ((total shuttled/total foraged)*100).

### Effort curve paradigm

Mice performed a within-subject experiment measuring effort modulation using the three different sized aperture faceplates of the nesting box (**Fig. 2C**). Mice underwent a foraging session under each of the three effort contingencies in a counterbalanced design (**S6**). Each session was 2 hours with at least a 2-day gap to avoid previous foraging experience impacting following sessions.

### Affective reactivity test

Mice underwent affective reactivity testing using the standard (**Fig.2A**) and larger (**Fig. 2D**) forage areas (**S7**). The ‘easy’ 1.5cm^2^ aperture faceplate was used for all tests. Mice underwent a foraging session under each of the conditions in a within-subject counterbalanced experimental design. 2-hour test sessions were run with at least a 2-day gap in between sessions to avoid previous experience impacting subsequent sessions.

### Novelty suppressed feeding test

To assess anxiety-related behaviour the novelty supressed feeding test was conducted. Mice were food restricted for 22 hours preceding task onset. Mice were run individually and in white light. Mice were placed into the NSFT arena (**S9**), containing a bowl of standard lab chow (Purina, UK) in the centre. Time the mice took to approach the bowl (latency to approach (s)) and eat the chow (latency to eat (s)) was recorded by the experimenter. If mice did not eat the chow within 10 minutes they were removed from the arena. Woodchip was redistributed and the chow bowl cleaned and food pellets replaced between mice. NSFT arena dimensions are provided in **S8**.

### Elevated plus maze

To assess anxiety-related behaviour outside of the context of food motivation, the elevated plus maze (EPM) test was conducted. Mice were placed on the EPM apparatus (**S10**) and left to explore for 5 minutes. A webcam positioned above the apparatus recorded activity and manual analysis of this footage was conducted using the in-house developed Novel Object Counter https://novel-object-video-counter.netlify.app/. Behaviours scored included number of entries into open and closed arms of the maze, time spent in the open arms and frequency of stretch attend posture (SAP). Mice were run individually, the apparatus was wiped down with 70% ethanol between runs, and all experiments took place in white light but during the active phase. EPM dimensions are provided in **S8**.

N=1 mouse was excluded from the study due to excessive stereotypic behaviour potentially driven by food restriction necessary for the NSFT.

### Operant training

Operant training was consistent with protocols reported in Marangoni et al (2023) and Jackson et al (2021). Training took place in 6 sound-proof operant boxes (Med Associates Inc.) (**Fig. 5**) and were run and recorded on Klimbic software (Conclusive Solutions Ltd., UK). Operant boxes consisted of two nose poke apertures either side of a magazine where reward pellets (20mg, rodent tablet, TestDiet, Sandown) were dispensed. Training consisted of 5 stages: magazine training, continuous reinforcement, fixed ratio (FR) 1, 2 and 4, outlined in **S11**. Mice were trained to complete trials, referring to the delivery of a reward pellet after completing the required number of nose pokes at the allocated aperture. Once all mice individually reached criteria for FR4 (two consecutive days of 30+ trials), where 4 nose pokes resulted in delivery of a reward, EfR testing began.

### Effort for Reward task

Mice underwent FR4 with the addition of a bowl of powdered lab chow (Purina, UK) positioned in front of the inactive aperture (**S12**). Chow was easily accessed via a 0.5 inch hole in the lid and layered with a metal mesh to prevent excess digging behaviour. Powdered chow was refilled to the same level and shaken to redistribute scent between mice.

During each 30-minute session, mice chose between the freely available but less palatable powdered chow, which represented a low effort, low value reward option or completed FR4 trials, which represented the high effort, high value reward option. Number of trials completed was recorded using Klimbic (Conclusive Solutions Ltd., UK) and weight of the chow bowl was measured before and after sessions. Cameras were positioned above the operant boxes with views of the three apertures to record activity at the chow bowl, and was analysed using the in-house designed Button Press Counter app https://github.com/dandovi/efr-video-analysis-tool. Videoed activities at the chow bowl are outlined in **S14**. EfR test sessions were preceded by a baseline session, see **S13**.

A further n = 4 mice did not complete the EfR test sessions due to development of excessive stereotypic behaviour.

### Statistical analysis

All data were tested for normality using a Shapiro-Wilk test. For single-factor analysis, if data were normally distributed, a repeated measure (RM) one-way ANOVA (Geisser-Greenhouse corrected) was used. Where a significant main effect was observed (p<0.05), Dunnett’s post hoc test was performed. If data violated normality, a Friedman test was used, and if significant, Dunn’s post hoc comparisons were performed. For two-factor analysis, a RM two-way ANOVA was used, and if significant effects occurred, Sidak’s post-hoc comparisons were applied. A paired t-test was used to compare two groups within subject. Outliers were defined as values outside ±2 standard deviations from the mean. Outliers/missing data points are provided in **S15**. Data were analysed in GraphPad Prism v 10.4.1.

## Results

### Effort modulation and foraging levels differed between mouse strains

Strains showed differences in initial arena exploration and performance in the Effort Based Forage task.

During three sessions of initial habituation to an empty arena, entries into the forage area decreased by session 3 in C57BL/6J mice but increased in 129sv and BALB/c mice (p<0.05). In the latter two sessions, BALB/c had the greatest number of entries compared to the other strains (p<0.001, **S16**).

In the effort curve paradigm, there was an aperture*strain interaction (F_(4,66)_=6.271, p=0.0002) and a main effect of aperture on total nesting material foraged (F_(1.904, 62.84)_=47.48, p<0.0001), where C57Bl/6J mice foraged more from the 1.5cm^2^ and 1cm^2^ apertures than the 0.75cm^2^ aperture (p=0.0003 and p=0.0003 respectively), and more from the 1.5cm^2^ than the 1cm^2^ aperture (p=0.0012). 129sv mice also foraged more with increasing aperture size, with higher foraging levels in 1.5cm^2^ and 1.0cm^2^ than 0.75cm^2^ (p=0.0001 and p=0.0478), and more in 1.5cm^2^ than 1.0cm^2^ (p=0.0478). BALB/c mice only foraged more with the 1.5cm^2^ aperture compared to the 0.75cm^2^ aperture (p=0.0277). There was also a main effect of strain (F_(2,33)_=35.60, p<0.0001), where C57Bl/6J and 129sv mice consistently foraged more than BALB/c mice from the 0.75cm^2^ aperture (p=0.0144 and p=0.0009 respectively), the 1cm^2^ aperture (p<0.0001 and p=0.0098 respectively), and the 1.5cm^2^ aperture (p=0.0011 and p<0.0001 respectively). 129sv mice foraged more than C57BL/6J mice at the 0.75cm^2^ aperture (p=0.0078) and 1.5cm^2^ aperture (p=0.0166) (**fig.3.A**).

In the total shuttled measure, there was a strain*aperture interaction (F_(4,66)_=4.502, p=0.0028), a main effect of strain (F_(2,33)_=17.64, p<0.0001), and of aperture (F_(1.960, 64.67)_= 22.53, p<0.0001). C57Bl/6J mice shuttled more material from the 1.5cm^2^ aperture compared to the smaller 1.0cm^2^ and 0.75cm^2^ apertures (p=0.0114 and p=0.0043 respectively). Both 129sv and BALB/c mice shuttled more material at the 1.5cm^2^ aperture than at the smallest 0.75cm^2^ aperture only (p=0.0027 and p=0.0487 respectively). C57Bl/6J and 129sv mice shuttled more material than BALB/c mice in the 0.75cm^2^ (p=0.0106 and p=0.0241 respectively) and 1.5cm^2^ (p=0.0061 and p=0.0002 respectively) apertures. At the moderate 1.0cm^2^ aperture, 129sv mice shuttled more than C57Bl/6J and BALB/c mice (p=0.0385 and p=0.0018 respectively) (**fig.3.B**).

In the % nesting material shuttled measure there was a main effect of strain (F_(2,33)_=6.302, p=0.0048) but not of aperture, nor any aperture*strain interaction. At the moderate 1.0cm^2^ aperture, 129sv mice displayed a higher % nesting material shuttled than both C57Bl/6J and BALB/c mice (p=0.0275 and p=0.0045 respectively). At the largest 1.5cm^2^ aperture, 129sv mice showed a higher % shuttled than BALB/c mice (p=0.0087) but not C57Bl/6J mice (**fig.3.C**).

### Affective reactivity measured in the EBF task differed between mouse strains

Comparing foraging levels in the standard forage area versus larger forage area, a main effect of strain (F_(2,33)_=28.24, p<0.0001), area size (F_(1,33)_=59.38, p<0.0001), and a strain*area interaction (F_(2,33)_=15.48, p<0.0001) on total nesting material foraged was observed. The larger forage area reduced total nesting material foraged by C57Bl/6J and 129sv mice (p=0.0001 and p<0.0001 respectively) but did not reduce foraging in the BALB/c mice. In the standard forage area condition, consistent with foraging behaviour in the effort curve paradigm, 129sv mice foraged more than C57Bl/6J and BALB/c mice (p<0.0001), and C57BL/6J mice foraged more than BALB/c mice (p=0.0009). In the large forage area condition, 129sv mice also foraged more than BALB/c mice (p=0.0388) and trended towards foraging more than C57Bl/6J mice (p=0.053) (**fig.3.D**).

There was also a main effect of strain (F_(2,33)_=7.499, p=0.0021), area size (F_(1,33)_=14.11, p=0.0007) and an area size*strain interaction (F_(2,33)_=4.358, p=0.0209) on total nesting material shuttled between standard and large forage areas. 129sv mice displayed a reduction in shuttling in the large area compared to the standard area (p=0.0007), and C57Bl/6J mice showed a trend towards the same reduction (p=0.0712). However, BALB/c mice showed no difference in shuttling behaviour between area sizes. In the standard area condition, 129sv mice shuttled more than C57Bl/6J and BALB/c mice (p=0.0234 and p<0.0001 respectively), and C57Bl/6J mice trended towards shuttling more than BALB/c mice (p=0.0934) (**fig.3.E**).

There were no main effects of strain, area size or a strain*area interaction on % nesting material shuttled (**fig.3.F**).

**Figure 3.**
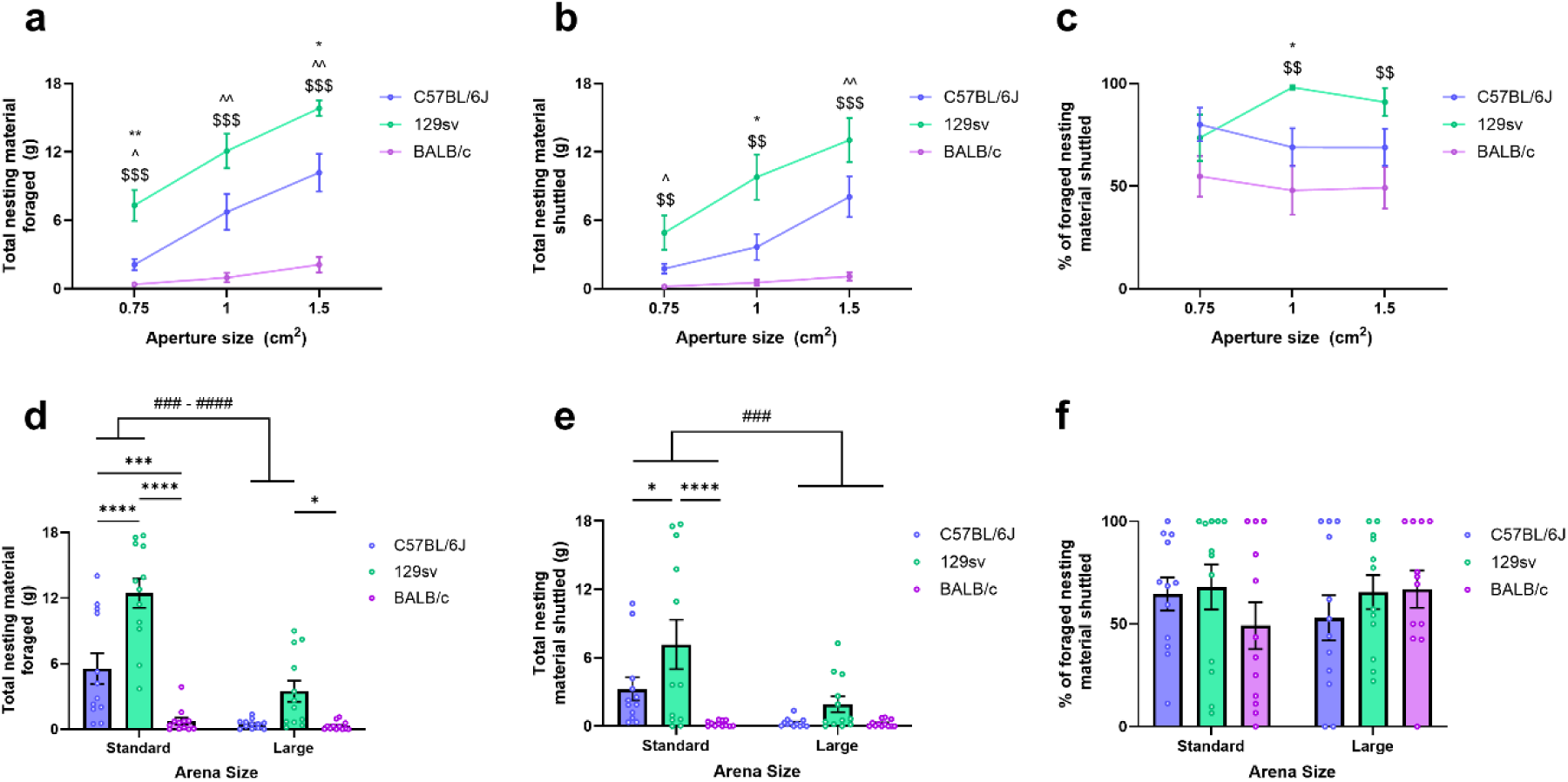
Strain differences in effort-based modulation and affective reactivity revealed in the EBF task. **A** 129sv mice foraged more nesting material than BALB/c mice across all aperture sizes (p<0.001) and more than C57Bl/cJJ mice at the 0.75 and 1.5cm^2^ apertures (p<0.01-0.05). C57Bl/6JJ mice foraged more than BALB/c mice across all aperture sizes (p<0.01-0.05). **B** 129sv mice shuttled more nesting material than BALB/c mice across all aperture sizes (p<0.001-p<0.01) and C57Bl/6JJ mice at the 1.0cm^2^ aperture (p<0.05). C57Bl/6JJ mice shuttled more than BALB/c mice at the 0.75cm^2^ and 1.5cm^2^ aperture (p<0.01-0.05). **C** 129sv mice had a greater % shuttled than BALB/c mice at the 1.0cm^2^ and 1.5cm^2^ apertures and C57Bl/6JJ mice at the 1.0cm^2^ aperture. **D** 129sv and C57Bl/6JJ mice show a reduction in foraging in the larger forage arena (p<0.001-p<0.0001). In the standard forage arena size, 129sv mice foraged more than C57Bl/6JJ and BALB/c mice, and C57Bl/6JJ mice foraged more than BALB/c mice (p<0.01-p<0.0001). In the larger forage arena, 129sv foraged more than BALB/c mice only (p<0.05). **E** 129sv and C57Bl/6JJ mice show a reduction in shuttling in the larger forage arena (p<0.001). In the standard forage arena size, 129sv mice shuttled more than C57Bl/6JJ and BALB/c mice (p<0.05-p<0.01). In the larger forage arena, there was no strain difference in shuttling. **F** There was no effect of arena size on % shuttled. Data are mean ± SEM with data points overlaid. Where data were non-normally distributed, median ± interquartile range were used instead. : - 12Ssv vs BALB/c, ^ - C57Bl/6JJ vs BALB/c, * - C57Bl/cJJ vs 12Ssv, # - within subject comparison.

### Extrinsic effort related choice behaviour differed between strains

Strains also showed differences in Effort for Reward task performance. There was a main effect of strain on number of high-effort/high-reward trials completed in the EfR task (F_(2,28)_=5.276, p=0.0114), with BALB/c mice completing more trials than 129sv mice (p=0.0082) and no differences between C57Bl/6J and 129sv or BALB/c mice (**fig.4.A**). There was also a main effect of strain on time spent at the chow bowl (F_(2,26)_=5.680, p=0.0090), with 129sv mice spending more time at the chow bowl than BALB/c mice (p=0.0095) (**fig.4.E**). Strain also affected the number of bouts at the chow bowl (F_(2,28)_=8.947, p=0.0010), with 129sv mice visiting the chow bowl more than C57Bl/6J and BALB/c mice (p=0.0097 and p=0.0013 respectively) (**fig.4.F**). There was no effect of strain on chow consumed, latency to complete first trial, or latency to approach the chow bowl (p>0.05) (**fig.4.B-D**).

Using foraging level data at the moderate aperture of the effort curve, simple linear regression showed no correlation between number of high effort, high reward trials completed in the EfR task and foraging level in the EBF task (**S17**).

**Figure 4.**
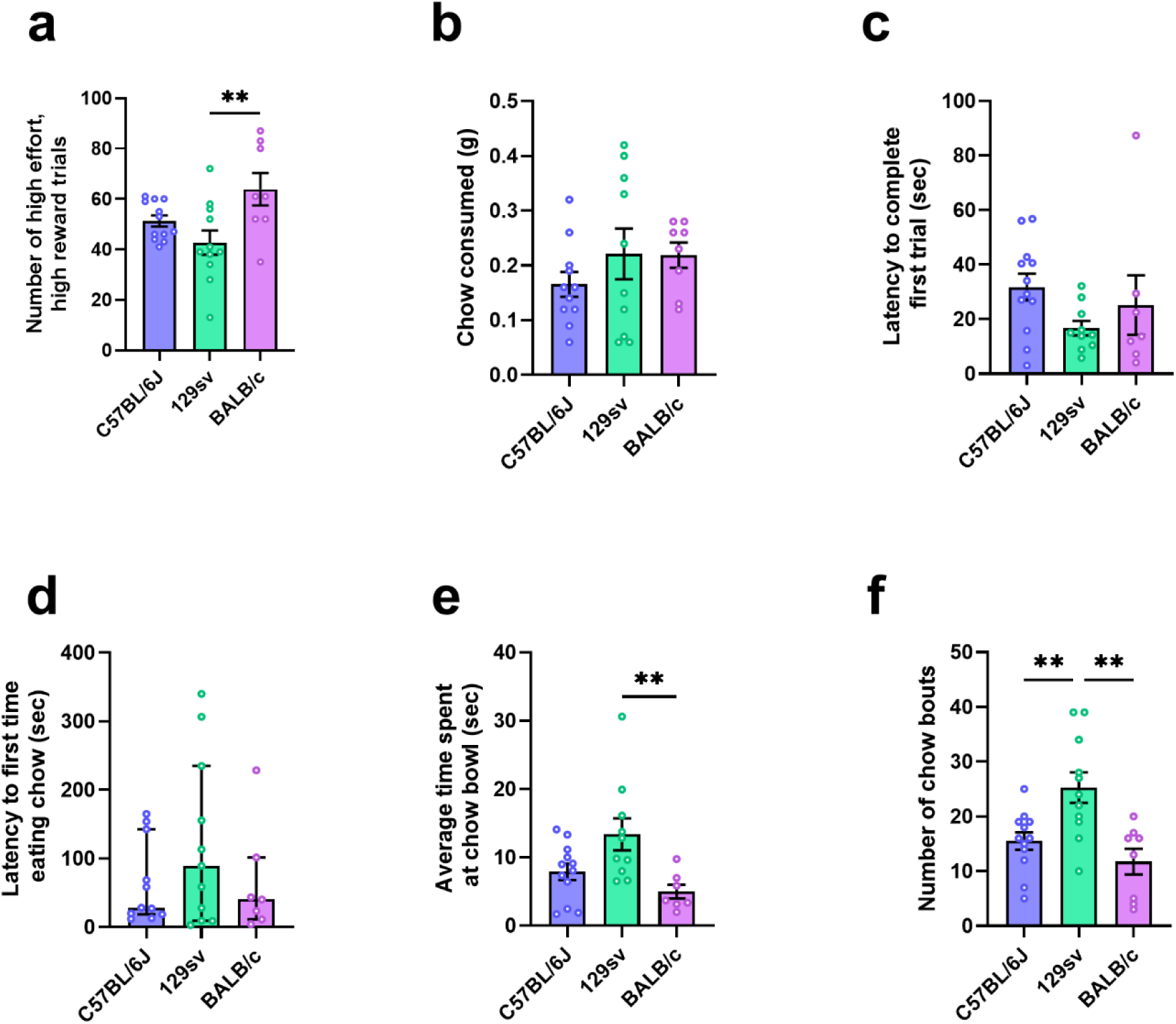
BALB/c mice showed a higher effort-based motivation in the EfR task. **A** BALB/c mice completed a higher number of trials than 129sv mice (p<0.01). **B** There was no strain difference in chow consumed, **C** latency to complete first trial or **D** latency to first time eating chow (p>0.05). **E** 129sv mice spent longer engaging with the chow bowl than BALB/c mice (p<0.01) and had a greater number of bouts at the chow bowl than both C57Bl/cJJ mice and BALB/c mice (p<0.01). * - strain comparison. Data are mean ± SEM with data points overlaid. Where data were non-normally distributed, median ± interquartile range were used instead.

### Strains display differing levels of anxiety-related behaviours

In the EPM, there was a main effect of strain on % time spent in the open arms (F_(2,31)_=3.985, p=0.0288) where C57Bl/6J mice spent more time in the open arms than 129sv mice (p=0.0231) (**fig.5.A**). There was also a main effect of strain on SAP frequency (F_(2,30)_=6.181, p=0.0057) where 129sv mice showed a higher count than BALB/c mice (p=0.0044) (**fig.5.B**). The number of arm entries was also affected by strain (F_(2,30)_=13.25, p<0.0001), where C57Bl/6J and 129sv mice displayed higher frequencies of entries than BALB/c mice (p<0.0001 and p=0.0290 respectively) (**fig.5.CsD**).

In the NSF test, there was no effect of strain on latency to approach the chow bowl, but there was a trend towards an effect of strain on latency to eat chow (F_(2, 31)_=3.129, p=0.0578) (**fig.5.EsF**).

There was no correlation between these anxiety relevant output measures and foraging in the EBF task, except for SAP counts in C57BL/6JJ mice where a weak positive correlation was observed (R^2^ = 0.4278, p=0.04278) (**S17**). All data are summarised in tables 1 and 2.

**Figure 5.**
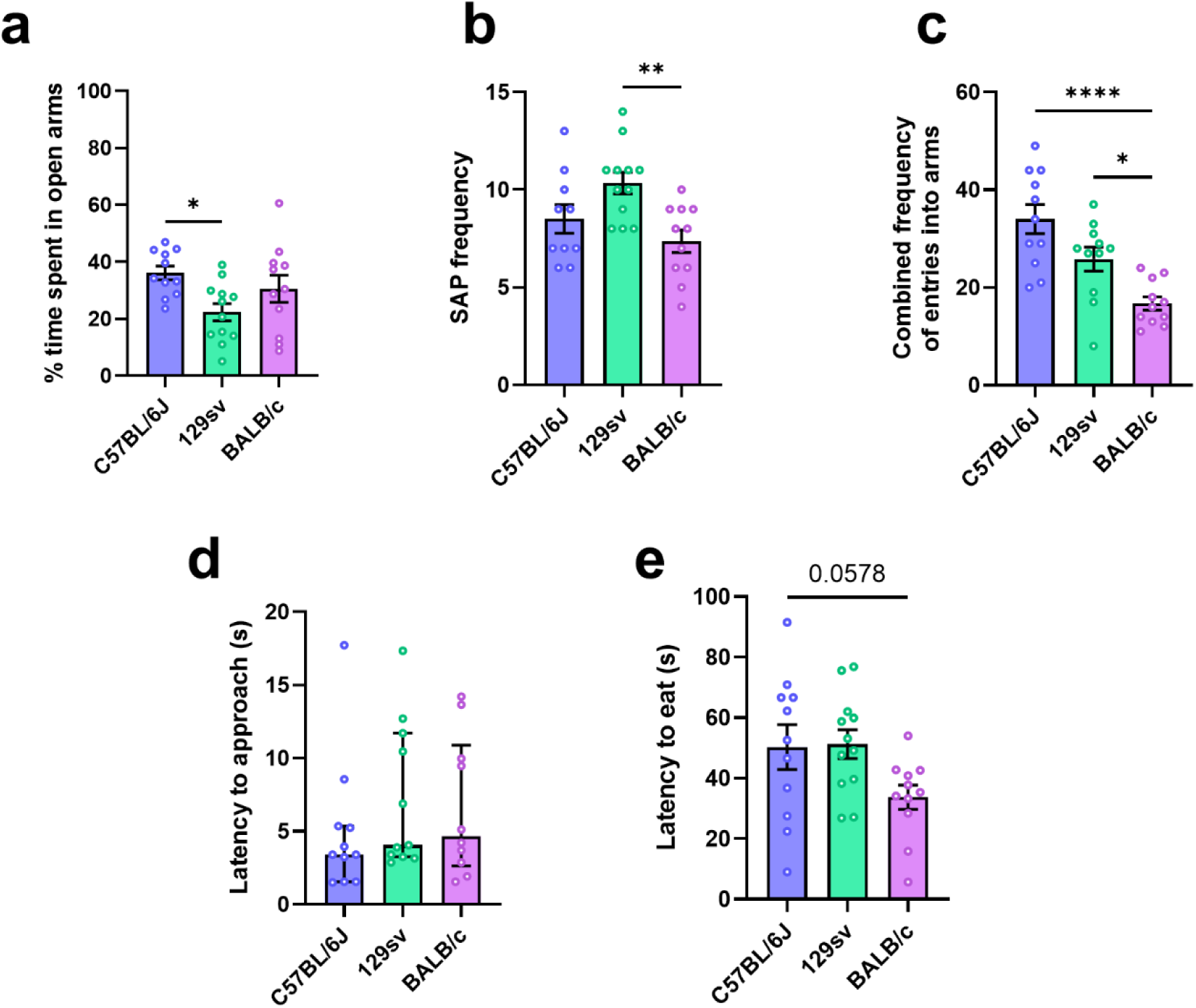
123sv mice display a higher anxiety-related phenotype compared to the other strains. **A** 129sv mice had a lower % time spent in the open arms of the EPM compared to C57Bl/cJJ mice (p<0.05). **B** 129sv mice had a greater SAP frequency than BALB/c mice (p<0.01). **C** BALB/c mice had a lower combined number of entries into the arms of the EPM compared to 129sv mice (p<0.05). 129sv had a lower combined number of entries than C57Bl/cJJ mice (p<0.0001). **D** There was no effect of strain on latency to approach the chow bowl in the NSFT. BALB/c mice trended towards a lower latency to eat in the NSFT compared to C57Bl/cJJ mice. * - strain comparison. Data are mean ± SEM with data points overlaid. Where data were non-normally distributed, median ± interquartile range were used instead.

**Table 1.** Data summary. EBF-effort based forage, EfR-effort for reward, EPM-elevated plus maze, NSFT-novelty supressed feeding test.

| Task | Measure | Strain comparisons |  |  |
| --- | --- | --- | --- | --- |
|  |  | C57Bl/6JJ | 129sv | BALB/c |
| Habituation to EBF task | Mean bouts/minute | = 129sv<br>< BALB/c | = C57Bl/6JJ<br>< BALB/c | > C57Bl/6JJ<br>> 129sv |
| Effort curve test in EBF task | Total foraged | < 129sv<br>> BALB/c | > C57Bl/6JJ<br>> BALB/c | < C57Bl/6JJ<br>< 129sv |
|  | Total shuttled | < 129sv<br>> BALB/c | > C57Bl/6JJ<br>> BALB/c | < C57Bl/6JJ<br>< 129sv |
|  | % shuttled | < 129sv<br>= BALB/c | > C57Bl/6JJ<br>> BALB/c | = C57Bl/6JJ<br>< 129sv |
| Affective reactivity test in EBF task | Total foraged | < 129sv<br>> BALB/c | > C57Bl/6JJ<br>> BALB/c | < C57Bl/6JJ<br>< 129sv |
|  | Total shuttled | < 129sv<br>= BALB/c | > C57Bl/6JJ<br>> BALB/c | = C57Bl/6JJ<br>< 129sv |
|  | % shuttled | = 129sv<br>= BALB/c | = C57Bl/6JJ<br>= BALB/c | = C57Bl/6JJ<br>= 129sv |
| NSFT | Latency to approach (s) | = 129sv<br>= BALB/c | = C57Bl/6JJ<br>= BALB/c | = C57Bl/6JJ<br>= 129sv |
|  | Latency to eat (s) | = 129sv<br>= BALB/c | = C57Bl/6JJ<br>= BALB/c | = C57Bl/6JJ<br>= 129sv |
| EPM | % time spent in open arms | > 129sv<br>= BALB/c | < C57Bl/6JJ<br>= BALB/c | = C57Bl/6JJ<br>= 129sv |
|  | SAP frequency | = 129sv<br>= BALB/c | = C57Bl/6JJ<br>> BALB/c | = C57Bl/6JJ<br>< 129sv |
|  | Combined frequency of entries into arms | = 129sv<br>> BALB/c | = C57Bl/6JJ<br>> BALB/c | < C57Bl/6JJ<br>< 129sv |
|  | Frequency of entries into open arms | > 129sv<br>> BALB/c | < C57Bl/6JJ<br>> BALB/c | < C57Bl/6JJ<br>< 129sv |
| EfR task | Chow consumed (g) | = 129sv<br>= BALB/c | = C57Bl/6JJ<br>= BALB/c | = C57Bl/6JJ<br>= 129sv |
|  | Trials completed | = 129sv<br>= BALB/c | = C57Bl/6JJ<br>< BALB/c | = C57Bl/6JJ<br>> 129sv |
|  | Latency to complete first trial (s) | = 129sv<br>= BALB/c | = C57Bl/6JJ<br>= BALB/c | = C57Bl/6JJ<br>= 129sv |
|  | Average time spent at chow bowl (s) | = 129sv<br>= BALB/c | = C57Bl/6JJ<br>> BALB/c | = C57Bl/6JJ<br>< 129sv |
|  | Number of chow bouts | < 129sv<br>= BALB/c | > C57Bl/6JJ<br>> BALB/c | = C57Bl/6JJ<br>< 129sv |
|  | Latency to first time eating chow (s) | = 129sv<br>= BALB/c | = C57Bl/6JJ<br>= BALB/c | = C57Bl/6JJ<br>= 129sv |

**Table 2.**
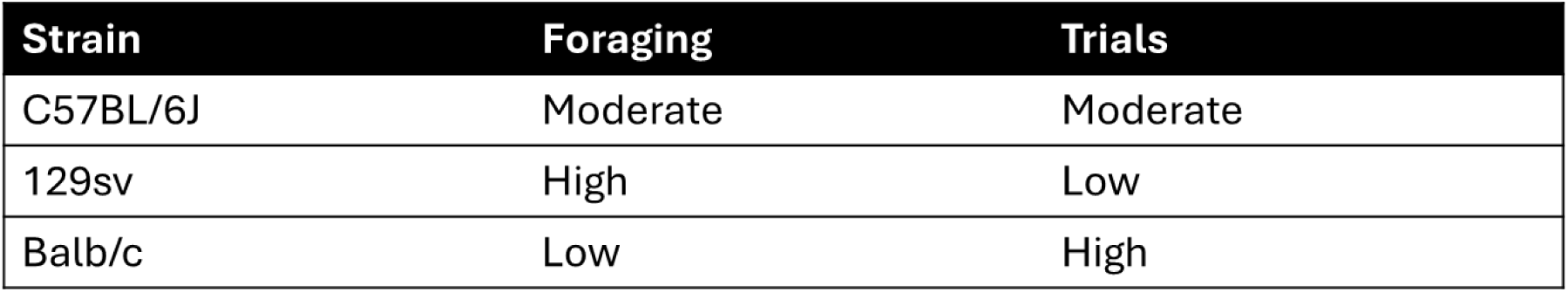
Low, moderate and high descriptors are based on relative performance between strains based on mean differences. Foraging refers to the total nesting material foraged output measure within the EBF task while trials refer to number of high effort, high reward trials completed in the operant EfR task.

## Discussion

Our findings reveal that different mouse strains exhibit distinct behavioural profiles across the two different motivation tasks which are not directly related to an anxiety-related phenotype. Specifically, we find that in the EBF task, 129sv mice show the highest overall level of foraging, while BALB/c mice showed the lowest. This indicates that 129sv mice are in a higher motivational state than BALB/c mice. In direct contrast, use of the operant EfR task showed that BALB/c mice completed higher numbers of high effort, high reward trials while 129sv mice showed higher levels of engagement with the low effort, low reward option. This indicates that BALB/c mice are in a higher motivational state than 129sv mice. Conventional tests of anxiety-relevant processes showed that 129sv mice had aspects of a higher anxiety-related phenotype than the other strains, indicating that the poor performance of BALB/c mice in the EBF task was not driven by anxiety-relevant behaviour. This divergence in findings across tasks further adds to our previous pharmacological studies which indicate that intrinsic and extrinsic motivation are differentially modulated.

The following discussion will consider how these findings contribute to our understanding of inter-strain variation in motivation phenotype and differences between intrinsic and extrinsic motivational processes.

### Strain differences reveal divergence between motivation tasks

The ability to adjust behaviour in response to changes in effort requirement is consistent with a healthy motivation system. The inability to appropriately adjust this behaviour, or disrupted sensitivity to effort requirement is indicative of motivation disruption common across many neurological and psychiatric disorders (Le Heron et al., 2018b, Mehrhof and Nord, 2025). Consistent with previous work utilising the same strain (Xeni et al., 2024, Xeni et al., 2025), C57Bl/6JJ mice altered their foraging behaviour relative to the effort required to forage, foraging more at the largest aperture relative to the smallest aperture. 129sv mice were similarly able to display this pattern of effort-based modulation of foraging behaviour. 129sv mice have been of scientific interest due to impairments observed in some aspects of working memory linked to a mutation in the *Disc1* gene relevant to Schizophrenia (Koike et al., 2006, Juan et al., 2014). Their performance in motivation-related tasks has not previously been examined. However, the present work suggests that intrinsic motivation-based processes and effort-based modulation is conserved. Conversely, BALB/c mice showed a lesser change in foraging level across aperture size which may be driven by a low overall foraging level, creating a floor effect. Consistent with this, BALB/c mice foraged less than 129sv mice across all aperture sizes and less than C57BL/6JJ mice at the smallest and largest aperture sizes. This comparatively low level of foraging indicates low intrinsic motivation in this strain.

As with 129sv mice, there is limited work on the specific assessment of motivation and/or goal-directed behaviour in BALB/c mice relative to other strains. However, multiple studies have shown that BALB/c mice show superior learning and higher levels of engagement in operant conditioning studies (Johnson et al., 2010, Johnson et al., 2009). These studies require the shaping of a specific behaviour to receive an external food reinforcer, and higher levels of responding are indicative of a higher overall motivational state. Consistent with this, the use of the Effort for Reward task in the present study showed that BALB/c mice completed more high effort, high reward trials and showed lower engagement with the low effort, low reward option compared to 129sv mice, indicative of a higher motivational state and contrasting EBF findings. This divergence in findings reveals important differences between task context and the nature of the motivational process engaged.

While both tasks utilise effort-based processes to assess motivational state, they differ considerably in external context. In particular, the EfR task requires a simplified environment, conditioning training to shape a behavioural response, and is reinforced by the delivery of palatable food. In contrast, the EBF draws on innate, untrained behavioural processes that are not guided by food reinforcement. BALB/c mice have been shown to have elevated impulsivity (Otobe and Makino, 2004) and perseverative behaviours (Zimmermann et al., 2016) compared to other strains that may favour responding in the EfR task environment where behaviours are externally shaped and directed but less favourable in a complex environment where unrestricted mice are not driven to perform one action over another. This aligns with the dissociable effects seen with certain drug treatments in the EBF versus the EfR task. The psychostimulant amphetamine increases food motivated operant task engagement by reducing the ability of the animal to shift between different behaviours within task (Hashemnia et al., 2020). Consistent with this, amphetamine improved performance in the EfR task, but was impairing in the EBF task (Xeni et al., 2024). As such, this work highlights that extrinsic and intrinsically motivated behaviours engage discrete motivational processes that can dramatically alter outcomes in different tasks and are also sensitive to phenotype.

Of note, this behavioural divergence across tasks has also been observed in a broader pharmacological context. It was found that acute administration of a range of serotonergic antidepressant drugs decreased foraging behaviour in the EBF task while increasing high effort, high reward trials in the EfR task (Xeni et al., 2025). This, together with the present findings indicates that partially distinct neurobiological mechanisms are engaged across intrinsic and extrinsically motivated tasks. These findings further support the case to move away from considering motivation on a single axis, as findings can diverge depending on the nature of the motivational process which has implications for phenotypic interpretation.

### Anxiety-related behaviours do not explain foraging deficits in the EBF task

The design of the EBF task environment with an enclosed home area and a tube leading to a more exposed forage area makes foraging output potentially sensitive to anxiety-related processes. Our present and previous work has shown that use of a larger and thus more aversive forage area supresses foraging behaviour, indicating that output is sensitive to environment-related conflict (Xeni et al., 2024). As such, the lower foraging levels observed in the BALB/c mice and higher levels in the 129sv mice have the potential to be attributed to changes in anxiety-related phenotype. Previous work examining anxiety-related levels across strains is mixed. For example, some work has found that BALB/c mice show lower levels of anxiety in the EPM compared to C57BL/6J mice (Avgustinovich et al., 2000), while others have found the opposite (Augustsson and Meyerson, 2004). Differences may depend on the age of the mouse and housing conditions among other factors, and as such, it was necessary to conduct measures of anxiety-related processing as part of the present work battery.

Anxiety-relevant processes specific to the EBF task environment can be examined using the EBF affective reactivity paradigm. An exaggerated or blunted change in foraging output measured in the standard versus large forage arena paradigm has been used to indicate changes in affective reactivity in disease models (Xeni et al., 2024). Here, BALB/c mice showed no change in foraging output between the standard and large forage areas. However, this is likely attributed to low foraging levels under standard conditions rather than an indication of affective blunting. C57BL/6JJ and 129sv mice showed an expected decrease in foraging in the aversive area, however % shuttled was unaffected. This contrasts with previous work where a larger forage area reduced % shuttled in C57BL/6JJ mice sourced from Envigo, UK, potentially highlighting sub-strain differences that warrant further exploration.

To further expand our assessment of anxiety-relevant behaviours between strains, conventional assays including the NSFT and the EPM were utilised. 129sv mice showed lower % time spent in the open arms of the EPM compared to C57BL/6JJ mice, and a greater SAP frequency compared to BALB/c mice. These measures suggest that 129sv mice had higher levels of anxiety-related behaviour compared to the other strains in the EPM. Of note, BALB/c mice showed the lowest movement in the anxiogenic EPM environment. This is consistent with previous work demonstrating lower levels of activity in an open field environment relative to other strains (Lipkind et al., 2004). This contrasts with the higher levels of activity observed in the forage area of the EBF task during habituation in the present work. This divergence in relative activity levels across the task environments suggests that BALB/c mice did not find the forage area anxiogenic. To account for limitations associated with the use of any one task, the NSFT was additionally utilised. Here, BALB/c mice trended towards a shorter latency to eat compared to C57BL/6JJ mice, indicating lower levels of anxiety-relevant behaviour. Overall, use of these conventional measures of anxiety-related behaviour indicates that 129sv mice show higher levels of anxiety-relevant behaviour while BALB/c mice show comparatively lower levels. Given that BALB/c mice showed the lowest level of foraging and 129sv the highest, this suggests that reduced foraging in the EBF task environment is not driven by elevated anxiety-related behaviour. This is supported by correlational analysis which found no relationship between anxiety measures and performance in the EBF task in these strains.

It is important to acknowledge that this work utilises male mice only. It is unclear how sex interacts with the observed strain differences in the context of motivation, however previous work has revealed sex and strain-mediated differences in social behavioural testing (Kopachev et al., 2022, An et al., 2011). Therefore, the addition of female mice could provide important insight into sex differences in cross-dimensional motivated behaviour. It is also important to note that differences in sub-strains were not examined. While three strains commonly used in neuroscience were selected, sub-strains may also demonstrate behavioural differences that warrant further exploration. For example, examination of behavioural differences between three C57BL/6J sub-strains revealed marked differences across a range of assays examining anxiety-related behaviour, memory and motor control (Matsuo et al., 2010) and the present work highlighted subtle foraging differences between C57BL/6J mice sourced from Janvier and Envigo.

## Conclusion

Here, we find that BALB/c mice show low levels of intrinsic motivation in the EBF task relative to the C57BL/6JJ and 129sv strains, but higher levels of extrinsic motivation in the EfR task. Their comparatively lower levels of anxiety-related behaviour suggest that their low EBF performance was driven by a specific motivational deficit. Our findings suggest that genetic background may shape motivational phenotype and importantly, that this phenotype can differ depending on the motivational process engaged. Notably, this work reveals that EBF and EfR task output diverges not just pharmacologically but also across strain phenotypes. This highlights the importance of testing both intrinsic and extrinsic motivation when phenotyping or assessing drug effect and provides further evidence for neurobiological differences between motivation domains.

## Acknowledgements

The authors would like to gratefully acknowledge the University of Bristol Animal Services Unit for care of the mice.

## Data availability

Data will become available on the Open Science Framework upon publication.

## References

3HS. 2024. The 3Hs Initiative: Housing, Handling and Habituation [Online]. Available: https://www.3hs-initiative.co.uk/ [Accessed 12/08/2024 2024].

An, X. L., Zou, J. X., Wu, R. Y., Yang, Y., Tai, F. D., Zeng, S. Y., Jia, R., Zhang, X., Liu, E. Q. C Broders, H. 2011. Strain and sex differences in anxiety-like and social behaviors in C57BL/6JJ and BALB/cJ mice. Exp Anim, 60, 111–23.

Augustsson, H. C Meyerson, B. J. 2004. Exploration and risk assessment: a comparative study of male house mice (Mus musculus musculus) and two laboratory strains. Physiology & Behavior, 81, 685–698.

Avgustinovich, D. F., Lipina, T. V., Bondar, N. P., Alekseyenko, O. V. C Kudryavtseva, N. N. 2000. Features of the Genetically Defined Anxiety in Mice. Behavior Genetics, 30, 101–109.

Cousins, M. S., Atherton, A., Turner, L. C Salamone, J. D. 1996. Nucleus accumbens dopamine depletions alter relative response allocation in a T-maze cost/benefit task. Behav Brain Res, 74, 189–97.

Davies, J., Jackson, M. G., Hinchcliffe, J. K., Mendl, M. C Robinson, E. S. J. 2025. Male mice prefer to live on their own. bioRxiv, 2025.05.02.651815.

Epstein, J. C Silbersweig, D. 2016. The Neuropsychiatric Spectrum of Motivational Disorders. Focus, 14, 499–509.

Fervaha, G., Foussias, G., Takeuchi, H., Agid, O. C Remington, G. 2016. Motivational deficits in major depressive disorder: Cross-sectional and longitudinal relationships with functional impairment and subjective well-being. Comprehensive Psychiatry, 66, 31–38.

Fervaha, G., Takeuchi, H., Lee, J., Foussias, G., Fletcher, P. J., Agid, O. C Remington, G. 2015. Antipsychotics and amotivation. Neuropsychopharmacology, 40, 1539–48.

Forstmeier, S. C Maercker, A. 2015. Motivational processes in mild cognitive impairment and Alzheimer’s disease: results from the Motivational Reserve in Alzheimer’s (MoReA) study. BMC Psychiatry, 15, 293.

Grayson, E. W., Xeni, F., Marangoni, C. C Jackson, M. G. 2025. The Effort-Based Forage Task: An Ethological Behavioral Test for Assessing Motivation and Apathy-Related Behavior in Mice. Curr Protoc, 5, e70195.

Hashemnia, S., Euston, D. R. C Gruber, A. J. 2020. Amphetamine reduces reward encoding and stabilizes neural dynamics in rat anterior cingulate cortex. Elife, 9.

Johnson, J. E., Pesek, E. F. C Christopher Newland, M. 2009. High-rate operant behavior in two mouse strains: A response-bout analysis. Behavioural Processes, 81, 309–315.

Johnson, J. M., Bailey, J. M., Johnson, J. E. C Newland, M. C. 2010. Performance of BALB/c and C57BL/6J mice under an incremental repeated acquisition of behavioral chains procedure. Behav Processes, 84, 705–14.

Juan, L.-W., Liao, C.-C., Lai, W.-S., Chang, C.-Y., Pei, J.-C., Wong, W.-R., Liu, C.-M., Hwu, H.-G. C Lee, L.-J. 2014. Phenotypic characterization of C57BL/6JJ mice carrying the Disc1 gene from the 129S6/SvEv strain. Brain Structure and Function, 219, 1417–1431.

Koike, H., Arguello, P. A., Kvajo, M., KArayiorgou, M. C Gogos, J. A. 2006. *Disc1* is mutated in the 129S6/SvEv strain and modulates working memory in mice. Proceedings of the National Academy of Sciences, 103, 3693–3697.

Kopachev, N., Netser, S. C Wagner, S. 2022. Sex-dependent features of social behavior differ between distinct laboratory mouse strains and their mixed offspring. iScience, 25, 103735.

Le Heron, C., Holroyd, C. B., Salamone, J. C Husain, M. 2018a. Brain mechanisms underlying apathy. *Journal of Neurology*, Neurosurgery & Psychiatry, 90, 302–312.

Le Heron, C., Plant, O., Manohar, S., Ang, Y.-S., Jackson, M., Lennox, G., Hu, M. T. C Husain, M. 2018b. Distinct effects of apathy and dopamine on effort-based decision-making in Parkinson’s disease. Brain, 141, 1455–1469.

LIpkind, D., Sakov, A., Kafkafi, N., Elmer, G. I., Benjamini, Y. C Golani, I. 2004. New replicable anxiety-related measures of wall vs. center behavior of mice in the open field. Journal of Applied Physiology, 97, 347–359.

Marangoni, C., Tam, M., Robinson, E. S. J. C Jackson, M. G. 2023. Pharmacological characterisation of the effort for reward task as a measure of motivation for reward in male mice. Psychopharmacology, 240, 2271–2284.

Matsuo, N., Takao, K., Nakanishi, K., Yamasaki, N., Tanda, K. C Miyakawa, T. 2010. Behavioral profiles of three C57BL/6J substrains. *Frontiers in Behavioral Neuroscience*, Volume 4–2010.

Mehrhof, S. Z. C Nord, C. L. 2025. A common alteration in effort-based decision-making in apathy, anhedonia, and late circadian rhythm. Elife, 13.

Morris, L. S., Grehl, M. M., Rutter, S. B., Mehta, M. C Westwater, M. L. 2022. On what motivates us: a detailed review of intrinsic v. extrinsic motivation. Psychol Med, 52, 1801–1816.

Muhammed, K., Manohar, S., Ben Yehuda, M., Chong, T. T. J., Tofaris, G., Lennox, G., Bogdanovic, M., Hu, M. C Husain, M. 2016. Reward sensitivity deficits modulated by dopamine are associated with apathy in Parkinson’s disease. Brain 139, 2706–2721.

Otobe, T. C Makino, J. 2004. Impulsive choice in inbred strains of mice. Behav Processes, 67, 19–26.

Rea, R., Carotenuto, A., Fasanaro, A. M., Traini, E. C Amenta, F. 2014. Apathy in Alzheimer’s Disease: Any Effective Treatment? The Scientific World Journal, 2014, 421385.

Salamone, J. D., Correa, M., Farrar, A. C Mingote, S. M. 2007. Effort-related functions of nucleus accumbens dopamine and associated forebrain circuits. Psychopharmacology (Berl), 191, 461–82.

Salamone, J. D., Steinpreis, R. E., Mccullough, L. D., Smith, P., Grebel, D. C Mahan, K. 1991. Haloperidol and nucleus accumbens dopamine depletion suppress lever pressing for food but increase free food consumption in a novel food choice procedure. Psychopharmacology, 104, 515–521.

Sansone, R. A. C Sansone, L. A. 2010. SSRI-Induced Indifference. Psychiatry (Edgmont), 7, 14–18.

Sultana, R., Ogundele, O. M. C Lee, C. C. 2019. Contrasting characteristic behaviours among common laboratory mouse strains. R Soc Open Sci, 6, 190574.

Xeni, F., Marangoni, C. C Jackson, M. G. 2024. Validation of a non-food or water motivated effort-based foraging task as a measure of motivational state in male mice. Neuropsychopharmacology.

Xeni, F., Marangoni, C., Lin, L., Robinson, E. S. J. C Jackson, M. G. 2025. Conditioned versus innate effort-based tasks reveal divergence in antidepressant effect on motivational state in male mice. Neuropsychopharmacology.

Zimmermann, K. S., Hsu, C.-C. C Gourley, S. L. 2016. Strain commonalities and differences in response-outcome decision making in mice. Neurobiology of Learning and Memory, 131, 101–108.

