## Supplementary document for "Male mouse strain variation reveals divergent phenotypes for extrinsic and intrinsic reward motivation"

**S1**

| Cohort | Stage/experiment | Age | Weight |
| --- | --- | --- | --- |
| EG1,2,3 (first half) | Arrival | 6 weeks | N/A |
|  | Handling habituation | 7 weeks | N/A |
|  | Habituation to EBF | 9 weeks | 22.8-26.8g |
|  | Effort curve | 14 weeks | 23.8-27.8g |
|  | Affective reactivity test | 16 weeks | 24.8-29.1g |
|  | NSFT | 18 weeks | 25.0-29.3g |
|  | EPM | 18 weeks | 26.8-30.8g |
|  | Operant training | 19 weeks | 24.7-30.3g |
|  | EfR | 24 weeks | 24.8-33.6g |
| EG4,5,6 (second half) | Arrival | 6 weeks | N/A |
|  | Handling habituation | 7 weeks | N/A |
|  | Habituation to EBF | 9 weeks | 23.7-26.6g |
|  | Effort curve | 11 weeks | 24.1-27.7g |
|  | Affective reactivity test | 12 weeks | 25.4-28.4g |
|  | NSFT | 14 weeks | 24.9-29.6g |
|  | EPM | 14 weeks | 26.0-29.9g |
|  | Operant training | 18 weeks | 26.5-29.7g |
|  | EfR | 22 weeks | 27.2-33.2g |

***S1. Summary of ages and weights of cohorts per stage or experiment.***

**S2**

Holding cages (open-top Techniplast 1284) contained a dome-shaped house, two cardboard tubes (one attached to the ceiling), a wooden labyrinth, nestlet, standard soft bedding material (IPS Product Supplies Ltd) and a layer of woodchip. Cages were cleaned once a week, and enrichment e.g. dome, tubes and labyrinth monitored for quality and renewed when necessary. Weights of mice were monitored at least once a week and maintained to at least 85% of their free-feeding weight relative to their normal growth curve.

**S3**
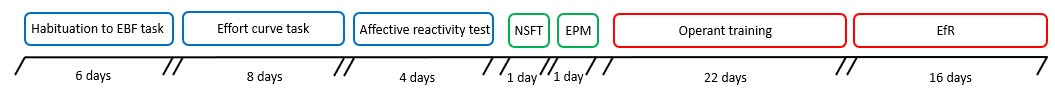

***S3.*** ***Experimental timeline.*** *Timeline illustrating the order experiments took place and the average duration (over two cohorts) in days of each. Blue boxes indicate the test was performed in the EBF arena, green boxes indicate assays measuring anxiety-related behaviour, and red boxes show assays conducted in operant boxes. EBF: effort-based forage, NSFT: novelty suppressed feeding test, EPM: elevated plus maze, EfR: effort-for-reward.*

**S4**

| Component | Dimensions |
| --- | --- |
| Home area | W 17.5cm, L 30.0cm, H 13.0cm |
| Home area lid | W 18.1cm, L 18.1 cm |
| Standard forage area | W 10.0cm, L 10.0cm, H 13.0cm |
| Large forage area | W 21.0cm, L 21.0cm, H 14.0cm |
| Connecting tube | L 21.0cm |
| Nesting material box body (3D printed) | W 3.5cm, L 8.0cm, H 10.0cm |
| Magnet bar (3D printed) | W 2.0cm, L 8.0cm, H 3.0cm |
| Nesting material box faceplate (3D printed) | W 0.3cm, L 7.8cm, H 10.0cm |
| Nesting material box lid (3D printed) | W 3.5cm, L 7.8cm, H 0.3cm |

***S4. Component dimensions for forage arena***

**S5**

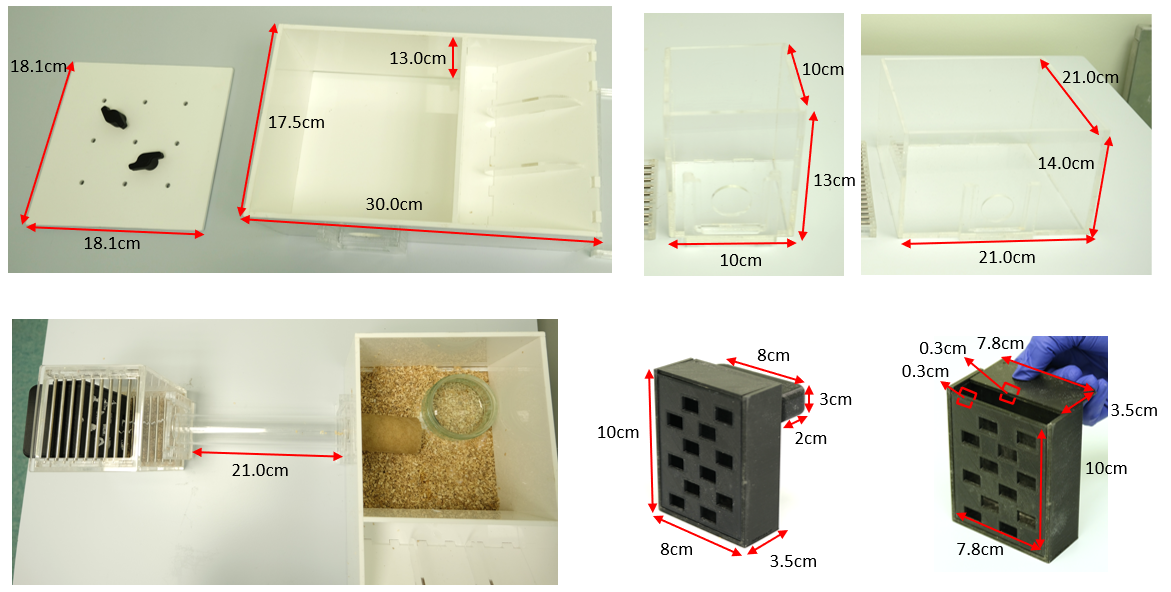

***S5. Components for forage arena labelled with dimensions.*** *From top left to bottom right; home area and home area lid, standard forage area, large forage area, connecting tube length, 3D printed nesting box (body and magnet dimensions), 3D printed nesting box (faceplate and lid dimensions).*

**S6**

|  | 3 x effort contingencies | | |
| --- | --- | --- | --- |
| Animal ID | Session 1 | Session 2 | Session 3 |
| Mouse 1 | Easy | Moderate | Difficult |
| Mouse 2 | Difficult | Easy | Moderate |
| Mouse 3 | Moderate | Difficult | Easy |

***S6. Example effort curve and affective reactivity experiment counterbalancing***

**S7**

|  | 2 x forage area sizes | |
| --- | --- | --- |
| Animal ID | Session 1 | Session 2 |
| Mouse 1 | Standard | Large |
| Mouse 2 | Large | Standard |

***S7. Example affective reactivity experiment counterbalancing***

**S8**

| Component | Dimensions |
| --- | --- |
| NSFT arena | W 39.7cm, L 39.7cm |
| NSFT chow bowl (distance from walls) | L 14.0cm |
| NSFT chow bowl | D 7.5cm, H 4cm |
| EPM arms (two closed) | L 78.2cm |
| EPM arm (one open) | L 35.5cm |
| EPM walls | H 15.5cm |
| EPM supports | H 60.0cm |

***S8. Component dimensions for NSFT and EPM apparatuses***

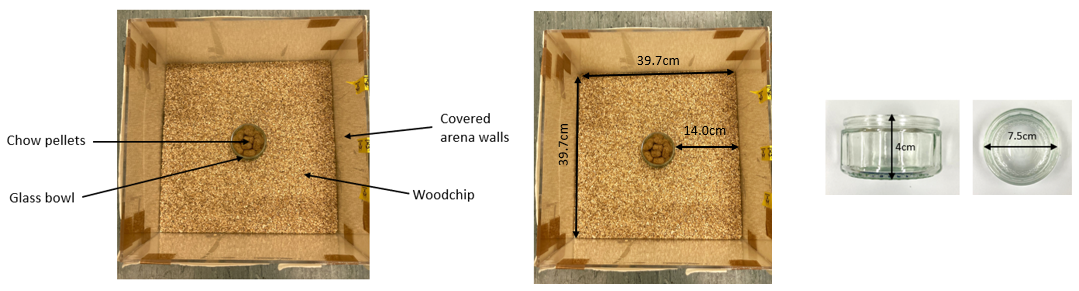
**S9**

***S9. NSFT apparatus and dimensions.*** *The NSFT arena consists of a woodchip-covered floor, covered walls to ensure mice cannot see outside the arena, and an open top. Standard lab chow pellets (Purina, UK) are placed into a glass bowl in the centre of the arena.*

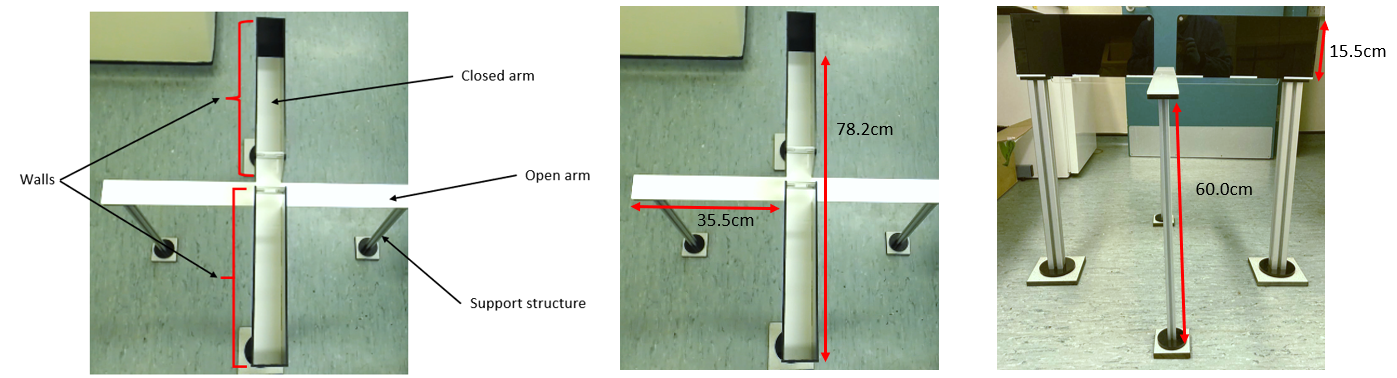
**S10**

***S10. EPM apparatus and dimensions.*** *The EPM includes two open arms (without walls) and two closed arms (with walls) and is secured with support structures. A camera is positioned directly above the apparatus to record activity.*

**S11**

| Stage | Criteria | Description |
| --- | --- | --- |
| Magazine training | 2 x sessions | Reward dispensed into magazine every 40 seconds |
| CRF | 5 x sessions | Reward dispensed if nose poke made in either aperture |
| FR1 | 2 consecutive sessions of 30+ trials | Reward dispensed if nose poke is made in allocated aperture |
| FR2 | 2 consecutive sessions of 30+ trials | Two nose pokes in allocated aperture required |
| FR4 | 2 consecutive sessions of 30+ trials | Four nose pokes in allocated aperture required |

***S11. Summary of operant training timeline, criteria and protocols***

**S12**

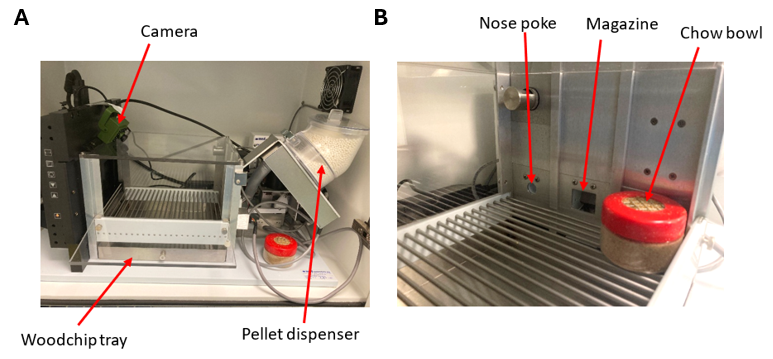

***S12. Operant box and EfR set-up.*** *Operant boxes (Med Associates Inc.) consisted of* ***A)*** *a tray covered with woodchip underneath the barred floor, a pellet dispenser connected to the magazine, and a camera positioned above the box with a view of the three apertures.* ***B)*** *The chow bowl has a 3cm diameter hole cut into the lid and covered with a metal mesh. It is positioned to cover the inactive nose poke (shown here as the right aperture).*

**S13**

| Cohort | Monday | Tuesday | Wednesday | Thursday | Friday |
| --- | --- | --- | --- | --- | --- |
| 1 | FR4 | FR4 | FR4 | FR4 | EfR |
|  | FR4 | EfR | Off | FR4 | EfR |
|  | FR4 | EfR | Off | FR4 | EfR |
| 2 | FR4 | EfR | Off | FR4 | EfR |
|  | FR4 | EfR | Off | FR4 | EfR |

***S13. Effort-for-reward experiment timetable***

**S14**

Activity during the EfR task was recorded with a camera and analysed using the Button Press Counter app, <https://github.com/dandovi/efr-video-analysis-tool>. The types of activity measured were latency to complete first trial, time spent at chow bowl, number of chow bouts (number of times mice visited the chow bowl), and latency to first time eating chow.

**S15**

| Experiment/analysis | Measure | Statistical exclusions in dataset |
| --- | --- | --- |
| Habituation to EBF task | Bouts/minute | N=2 outliers (1 C57Bl/6, 1 BALB/c) |
| Effort curve test | Total foraged | N=4 outliers (1 C57Bl/6, 1 129sv, 2 BALB/c) |
|  | Total shuttled | N=5 outliers (2 C47Bl/6, 3 BALB/c) |
|  | % shuttled | N=3 outliers (1 C57Bl/6, 2 129sv) |
| Affective reactivity test | Total foraged | N=4 outliers (1 C57Bl/6, 1 129sv, 2 BALB/c) |
|  | Total shuttled | N=4 outliers (2 C57Bl/6, 1 129sv, 1 BALB/c) |
|  | % shuttled | No exclusions |
| NSFT | Latency to approach | N=4 outliers (1 C57Bl/6, 1 129sv, 2 BALB/c) |
|  | Latency to eat | N=2 outliers (1 C57Bl/6, 1 BALB/c) |
| EPM | % time spent in open arms | N=2 exclusions; N=1 C57Bl/6 due to camera fault; N=1 BALB/c due to stereotypy |
|  | SAP | N=3 exclusions; N=1 C57Bl/6 due to camera fault; N=1 BALB/c due to stereotypy; N=1 outlier (C57Bl/6) |
|  | Combined entries | N=3 exclusions; N=1 C57Bl/6 due to camera fault; N=1 BALB/c due to stereotypy; N=1 outlier (129sv) |
|  | Entries into open arms | N=3 exclusions; N=1 C57Bl/6 due to camera fault; N=1 BALB/c due to stereotypy; N=1 outlier (129sv) |
| EfR test | Chow consumed | N=7 exclusions; N=5 (4 BALB/c, 1 129sv) due to stereotypy; N=2 outliers (1 C57Bl/6, 1 129sv) |
|  | Trials completed | N=5 exclusions; (4 BALB/c, 1 129sv) due to stereotypy |
|  | Latency to complete first trial | N=7 exclusions; N=5 (4 BALB/c, 1 129sv) due to stereotypy; N=2 outliers (1 129sv, 1 BALB/c) |
|  | Average time spent at chow bowl | N=7 exclusions; N=5 (4 BALB/c, 1 129sv) due to stereotypy; N=2 outliers (1 129sv, 1 BALB/c) |
|  | Number of chow bouts | N=5 exclusions (4 BALB/c, 1 129sv) due to stereotypy |
|  | Latency to first time eating chow | N=7 exclusions; N=5 (4 BALB/c, 1 129sv) due to stereotypy; N=2 outliers (1 C57Bl/6, 1 BALB/c) |
| Correlational analyses | Bouts/minute in EBF task | N=2 outliers |
|  | Total shuttled in EBF task | N=2 outliers |
|  | % time spent in open arms of EPM | N=1 outlier |
|  | Combined entries in EPM | N=1 outlier |
|  | Trials completed in EfR | N=2 outliers |
|  | All other measures | No exclusions |

***S15. Summary of data point outliers and exclusions per experiment or analysis***

**S16**

There was an interaction between strain and session (F_(3.912,64.54)_ = 11.27, p<0.0001), an effect of strain (F_(2,33)_ = 33.82, p<0.0001) but no effect of session. In the first session, 129sv mice had the lowest bouts/min compared to C57Bl/6J (p<0.0001) and BALB/c mice (p=0.0001) with no difference between C57Bl/6J and BALB/c strains. In the second session, BALB/c mice had a greater bouts/min than both C57Bl/6J (p=0.0004) and 129sv mice (p<0.0001), with no difference between C57Bl/6J and 129sv mice. In the final session, BALB/c mice again had greater bouts/min than both C57Bl/6J (p=0.0007) and 129sv mice (p<0.0001), with no difference between C57Bl/6J and 129sv mice.

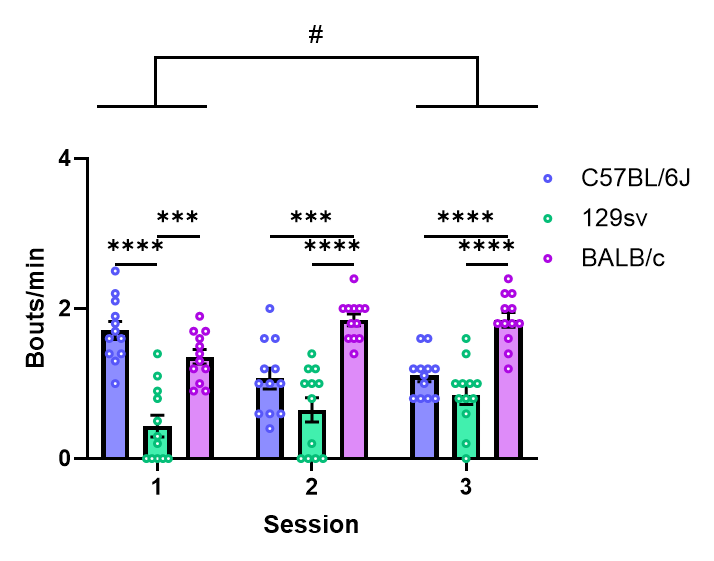

***S16.*** ***Strains differed in activity during habituation to EBF arena.*** *Number of entries into the empty forage area were counted over three sessions and normalised to time. 129sv mice showed the lowest bouts/min in the first session, and BALB/c mice displayed the greatest bouts/min in the latter two sessions.*

**S17**

There were no correlations between trials completed and total foraged in C57Bl/6J, 129sv or BALB/c strains. There was a weak positive correlation between SAP frequency and total foraged in C57Bl/6J mice (F(1,8) = 5.981, p = 0.0402), but no other correlations between total foraged and anxiety-relevant measures in C57Bl/6J, 129sv or BALB/c mice.

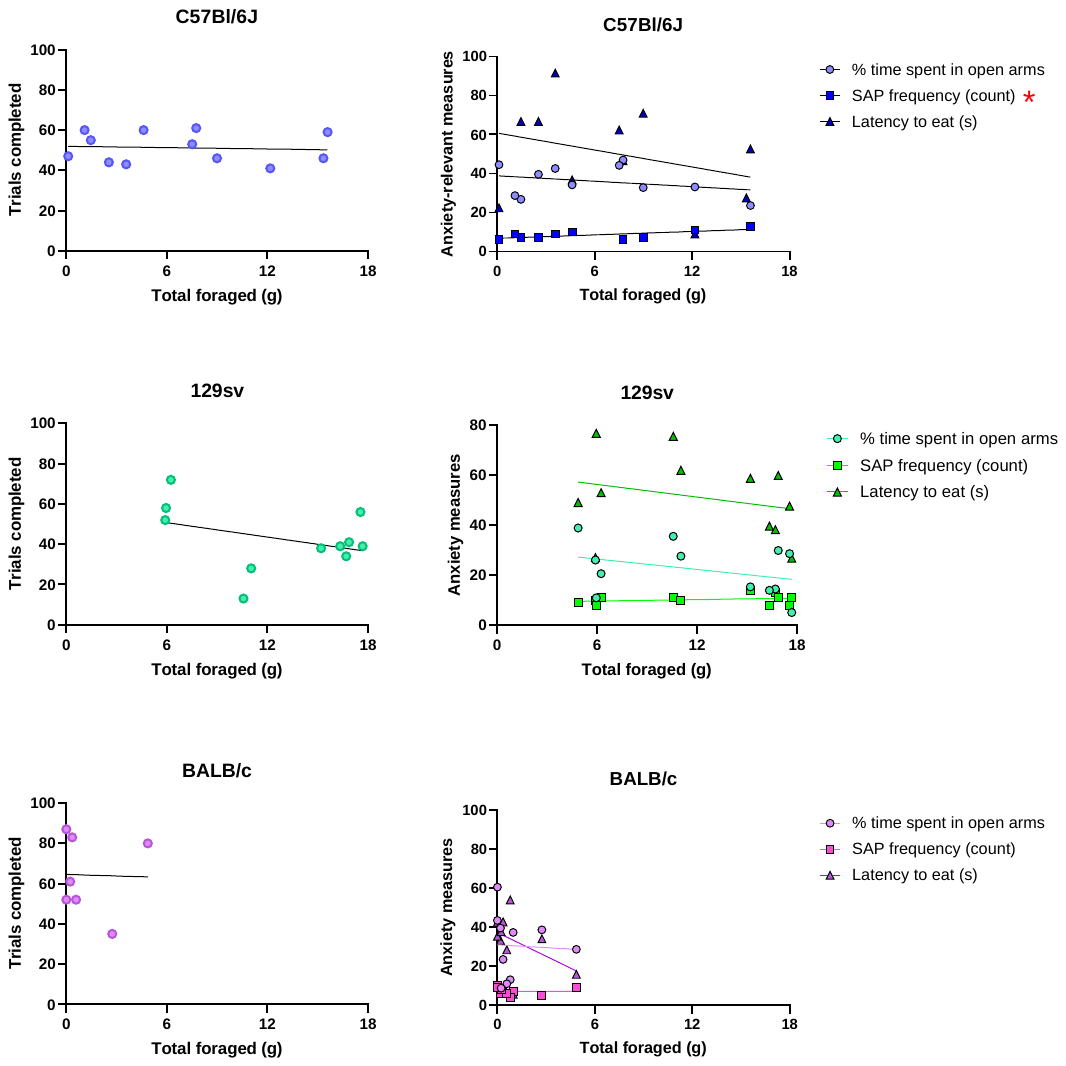

***S17. Correlational analyses show no relationship between trials completed and foraging and only a weak correlation between foraging and SAP frequency in C57Bl/6J mice.***
